# Reliability of tactile perception and suppression measurements

**DOI:** 10.1101/2024.08.20.608823

**Authors:** Dimitris Voudouris, Petros Georgiadis, Katja Fiehler, Belkis Ezgi Arikan

**Affiliations:** Experimental Psychology, Justus Liebig University Giessen, Germany; Department of Biology, York University, Toronto, Canada; Department of Neurology, Center for Translational Neuro- and Behavioral Sciences, University Hospital Essen and Experimental Psychology, Justus Liebig University Giessen, Germany

**Keywords:** detection thresholds, tactile adaptation, tactile gating, temporal effects

## Abstract

Tactile signals arising on one’s own body allow estimation of one’s own sensory state and foster interactions with the environment. However, tactile perception can be influenced by various factors. For instance, tactile perception is suppressed on a moving limb compared to when it is resting, a phenomenon termed tactile suppression. Here we examine whether tactile perception during resting and during movement is robust over shorter and longer time intervals. Participants had to detect tactile stimuli of various intensities on their index finger while this finger was resting or moving (finger extension). This detection task was performed on four sessions at separate days across a period of one month. We found that tactile perception during resting is robust within single sessions and across days. However, tactile perception during movement changed across days, but these changes lacked a clear systematic pattern. We further show that temporal changes in perception alone cannot fully account for the previously reported tactile suppression effects. Finally, split-half correlations reveal high consistency in the estimated perceptual measures, demonstrating that estimates of tactile perception are robust across measurement points.

## I. Introduction

Tactile signals provide humans with the ability to feel and successfully interact with their surroundings as these signals convey information about the properties of the environment and of one’s own movements (for a review, see [1]). For example, humans can use signals from their touch receptors to identify properties of surfaces and objects with which they interact, such as temperature [2], friction [3], edges [4] or shape [5]. In addition to perceiving the surrounding world, tactile signals are important for motor behavior, for instance by fostering the sense of body position [6] and by detecting object slippage during grasping [1]. Considering these, it may appear as a paradox that tactile sensitivity is suppressed on body parts just before and during voluntary movement [7-10]. This is reflected in the reduced perception of tactile stimuli when these are presented on a moving compared to a static limb.

This phenomenon in tactile perception, called tactile suppression, is evident in various human actions, such as finger abduction [11] and lifting [12], stroking movements [13], reaching [10], grasping [8], and even during whole-body tasks such as balance retention [14]. Suppression is thought to result from a feedforward-feedback mechanism that integrates priors about the underlying dynamics with sensory feedback to release a motor command, through which the brain estimates future sensory states of the body [15]. This estimate is compared with the actual sensory feedback and signals that match the prediction are suppressed. Consequently, just preparing to execute a voluntary movement can lead to tactile suppression on the body part that is planned to move, even if it eventually does not move [12, 16]. Likewise, tactile suppression is stronger during active than passive movements [17], likely because the latter are assumed to lack a feedforward motor command and thus predictions about the sensory states of the passively moved limb. However, passive movements do lead to tactile suppression, albeit to a lesser extent than active ones [17], which is taken as evidence that other, non-predictive, mechanisms may also be involved in this phenomenon [18]. Such mechanisms may refer to the masking of tactile probes by afferent signals [11], cognitive processes influencing the perception of the tactile probe [19, 20], or possibly other mechanisms, such as adaptation to tactile stimulation over time [21].

One of the most common ways to assess tactile perception in general, and tactile suppression specifically, is to conduct psychophysical experiments and estimate tactile detection [22, 23] or discrimination [10, 24] thresholds, or tactile sensitivity using signal detection measures [8, 17]. Typically, tactile perception is quantified while participants keep their stimulated body part at rest, and the effects of motor processes on perception are examined while participants perform instructed movements with the stimulated body part (e.g., [22, 23]). The resting and moving conditions are often presented in separate, counterbalanced blocks of several trials (e.g., [22, 23]), or in a single block with resting and moving trials being presented randomly and interleaved (e.g., [17]), with each of those blocks typically lasting up to 60 minutes. Sometimes, for instance when the experimental blocks last longer, the experiments are even split over separate days to keep each experimental session shorter (e.g., [17, 25]). In either case, experimenters estimate tactile perception in each of the resting and moving conditions separately and subsequently assess the effect of movement on tactile perception.

Repetitive exposure to sensory stimuli can have consequences on perception itself. These consequences can be observed as enhancement or decline of perceptual performance that may arise from perceptual learning or sensory adaptation, respectively. Specifically, repeated exposure or training can lead to sharpened perceptual performance (for a review see [26]), which can manifest in improved tactile detection. This is evident in studies where discrimination of auditory frequencies and of visual motion directions constantly improves with more exposure to the task, at least within a period of 7-8 sessions performed on consecutive days [27, 28], and if participants are exposed to a minimal number of trials [27]. Meanwhile, repeated exposure to a stimulus can sometimes lead to adaptation of the activated tactile receptors and the underlying neural pathways. This adaptation can result in an exponential decrease in sensory sensitivity, which can be reflected in poorer tactile detection after various intervals of suprathreshold stimulation [21, 29]. Perceptual learning and tactile adaptation can have implications in the assessment of tactile perception and tactile suppression. In the context of perceptual learning, tactile sensitivity may improve with more exposure to the stimuli or to the performed perceptual task, and thus perception may be better at the end of an experiment. Conversely, in the context of adaptation, tactile sensitivity may decrease with more exposure to stimuli, and thus perception may be poorer toward the end of an experiment. In other words, estimates of tactile perception and suppression may not be driven completely by the reported sensorimotor or cognitive factors (e.g., [8, 13]), but these estimates might be confounded by temporal changes in tactile sensitivity over the course of an experiment. Some studies explored this possibility by testing potential differences in tactile perception between early and late parts of the experimental procedure, with the comparisons typically providing no evidence for differences in tactile performance during resting, at least when the tested periods were within ∼30 min [30] and ∼90 min [31]. To the best of our knowledge, however, no study has assessed the reliability of tactile perception and suppression over various measurement points that take place within and across days. This is important because the reported effects of tactile suppression may be affected by such temporal effects that might require re-interpretation of previous results.

We conduct this study with three main aims. First, we investigate whether there is any change in tactile perception and suppression within and across experimental sessions that take place within one month. Second, we quantify the reliability of tactile perception and suppression measurements over the same period by quantifying how robust tactile perception and suppression measures are at different measurement points. Third, we connect the obtained findings with previous related work to examine whether the tactile suppression effects reported in the literature reflect a byproduct of temporal changes in perception, for instance, due to perceptual learning or tactile adaptation (see *Supplementary Material*). We asked healthy young human participants to perform a tactile detection task in which tactile stimuli were delivered to their index finger while their stimulated hand was resting or actively performing a finger extension. This task and the tactile stimuli are similar to those implemented in previous studies examining tactile suppression during hand movements (see *Supplementary Material*). The experiment consisted of four 60-min sessions performed on four different sessions over the period of one month. Each session comprised 12 mini-blocks; six mini-blocks with the finger resting and another six mini-blocks with the finger moving. We assessed tactile perception in each of these mini-blocks for each session separately. If tactile perception is influenced by perceptual learning or adaptation processes, we expect to see improved or worse sensitivity over time, respectively.

## II. Methods

### A. Participants

Twenty-six healthy adults participated in the study, but due to technical issues (i.e. failure in saving one datafile), our final sample comprised 24 adults (18 females, age range: 19-33 years; 6 males, age range: 20-35; mean age: 21.8 years old). This sample size is consistent with that of most other studies that investigated and found tactile suppression (for details, see *Supplementary Material*). All participants were recruited from pools of the Justus Liebig University Giessen and were right-handed according to the German translation of the Edinburgh Handedness Inventory ([32]; mean EHI score: 91; EHI range: 69-100). None of them had any known musculoskeletal or neurological issue that could influence their participation. After providing a detailed explanation of the study procedure, written consent was obtained from all participants. The study was approved by the local ethics committee of Justus Liebig University Giessen and was conducted in accordance with the Declaration of Helsinki (2013; except for §35, pre-registration). Participants received either 8€/hour or course credits for their efforts.

### B. Apparatus and setup

Participants sat in front of a table with their right index finger resting on a button box and their left hand positioned on a response keyboard. Brief (50 ms, 250 Hz) vibrotactile stimuli varying in intensity (peak-to-peak amplitude of 6, 12, 19, 25, 31, 38 μm) were transmitted using a custom-made vibrotactile device (Engineer Acoustics Inc., USA) to the dorsal part of the right index finger’s proximal phalanx. The vibration frequency, range and site of stimulus presentation were chosen to match the properties of stimuli used in previous studies that examined tactile suppression during hand movements (see *Supplementary Material*). This allowed us to relate any temporal changes in perception to previous results on tactile suppression. An Optotrak Certus system (NDI, Canada) recorded the position of an infrared marker fixed to the participant’s right index fingernail at 100 Hz.

### C. Procedure

Participants attended four 60-min measurement sessions, each on a separate day, following a specific timetable: the second session occurred the day following the initial measurement, the third session took place seven days after the first measurement, and the fourth session was performed thirty days after the initial session. The reason for recording perception on 60-min sessions is because most studies on tactile suppression have a similar duration (e.g., [10, 20, 23]). Moreover, we recorded on separate days because researchers sometimes split an experiment over separate days (e.g., [17, 25]). Although we could have measured perception in many other combinations of days, we chose these intervals as a convention to characterize possible changes over these measurement points. We took care that each participant’s four sessions were scheduled at roughly the same time of day.

Upon arrival at the lab, participants were given time to settle into a comfortable seated position. Their left hand rested on a table ∼30 cm in front of them and 20 cm to the left of their midline, positioned on the response keyboard. Their right hand rested on the button box ∼30 cm in front of them and 20 cm to their right of their midline. Hand placement was customized for individual comfort, considering variations in arm, hand, and finger lengths. Participants performed 12 mini-blocks, each containing one of two possible conditions: a resting (no-movement) and a movement condition. These conditions were presented in pairs of two consecutive mini-blocks, but the order of these conditions was randomized across both measurement days and participants. Each mini-block consisted of 48 trials. This allowed us to compare our results with previous work that examined and found tactile suppression with a similar number of trials (e.g., [22, 23, 30]). We ensured that the perceptual estimates were accurate as one can inspect in the individual psychometric functions (https://osf.io/wcrvd). Within each mini-block, each of the six possible vibrotactile stimuli was presented six times, and each of the no-stimulation (catch) trials was presented twelve times (20% of all trials), all in a random order. Figure 1 depicts the setup and the distribution of the measurement sessions.

**Fig. 1.**
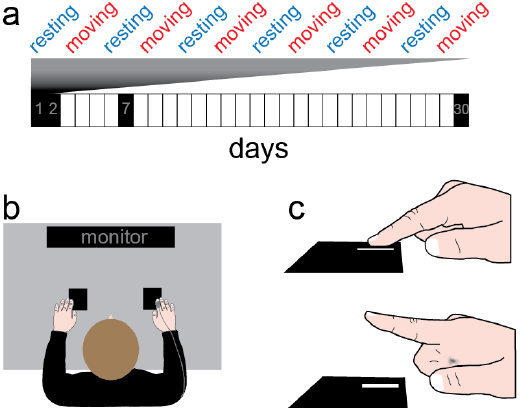
Setup and timeline. (a) Timeline of the experimental procedure. Each block indicates a calendar day, and black boxes indicate the measurement days. Each measurement day included 12 mini-blocks, six with resting and another six with moving trials. (b) Top view depiction of the setup. (c) Right index finger configuration in the static (upper) and moving (lower) trials.

In the resting condition, trial onset was defined by a key press with the left hand while the right index finger rested on the button box. The tactile stimulus was delivered to the right index finger between 210 and 280 ms (average: 245 ms) after the left hand pressed the key. Conversely, in the movement condition, participants pressed and held down the button box with their right index finger for at least 200 ms. Subsequently, they had to release the button by extending (i.e., moving up) their right index finger at the moment of their choice in a comfortable pace. In the moving condition, the stimulus on the right index finger occurred between 10 and 80 ms (average: 45 ms) after movement onset, so at the very minimum 210 ms after button press (i.e. trial onset). At the end of each trial, a question appeared on the monitor in front of the participants, asking them whether they detected a tactile stimulus or not. They responded to that question by pressing with any finger of their left hand one of two predefined keys on the response keyboard. There was no time limit for their responses. Each mini-block lasted ∼3 minutes, with a 2-minute break between mini-blocks, resulting in a duration of ∼60 minutes for each session (day). This represents the typical duration of several tactile suppression studies (e.g., [22, 23, 30, 33]).

### D. Data analysis

The first step in our analysis was to estimate detection thresholds for each participant. We did so separately for each mini-block and movement condition (i.e., resting and moving). Specifically, we fitted the detection responses with a cumulative normal distribution using *psignifit 4* [34] in Matlab 2023b (The Mathworks Inc., Natick, Massachusetts). We defined the detection threshold as the stimulus intensity at 50% correct of each resulting psychometric function. We did so separately for each of the six resting and moving mini-blocks within each session, obtaining six resting and six moving tactile detection thresholds per participant and session. Because we presented the resting and moving mini-blocks in an alternating order, we quantified tactile suppression by subtracting the detection threshold from a given resting mini-block from the detection threshold of the corresponding moving mini-block. This resulted in six values (*Δthreshold*) per participant and session that represent the effect of movement on tactile perception. Specifically, positive values indicate higher detection thresholds during movement than rest trials, reflecting higher movement-related tactile suppression.

Our first aim was to examine whether tactile perception and suppression changed within and across sessions (days). We implemented linear mixed models (LMMs) using the *lmerTest* package [35] in R Studio version 2023 4.2.3 [36]. The LMMs were fitted using the restricted maximum likelihood estimation. We initially checked model assumptions on a simple model consisting of fixed effects of movement, mini-block, day and random effect of participant with the *check_model* function. Visual inspection of model assumptions did not reveal any obvious deviations from normality of residuals, random effects, linear relationship, homogeneity of variance and multicollinearity. Two LMMs were conducted. The first LMM tested the effect of movement within and across days on the detection thresholds and included the fixed effects terms movement, mini-block, day as well as their interactions. To account for other (known) sources of variability, the model included the following random effects: a random intercept for participants, which accounts for overall within-subject variability and a random slope for movement, which accounts for the variability across participants associated as a function of movement. The results of this LMM are presented in Table 1. The second LMM examined the tactile suppression effect, i.e., Δthresholds, within and across days. In addition, as tactile suppression can be modulated by movement speed [37], we entered each participant’s average movement speed in each condition as a covariate of no interest. This LMM included the fixed effects terms mini-block, day and their interaction, average movement speed as covariate and the random effect term of participant. The results of this LMM are presented in Table 2. For all fixed effects of interest, we used dummy coding. We estimated the fixed effect terms with the *anova* function, which performs an F-test on the fixed effects using the Satterthwaite approximation [38]. For post-hoc pairwise comparisons and estimated marginal means, we used the *emmeans* package [39]. For multiple comparisons, we used Bonferroni adjustment, when necessary. Effect sizes were calculated using the *effectsize* package in R as Cohen’s *d* [40].

**TABLE 1.** STATISTICAL RESULTS FOR RAW THRESHOLDS.

|  | *SS* | *MS* | *dfs* | *F* | *p* |
| --- | --- | --- | --- | --- | --- |
| Movement | 5.9 | 5.9 | 1, 23 | 14.7 | < 0.001 |
| Mini-block | 20.3 | 4.1 | 5, 1058 | 10.1 | < 0.001 |
| Day | 9.1 | 3.1 | 3, 1058 | 7.6 | < 0.001 |
| Movement * mini-block | 8.1 | 1.6 | 5, 1058 | 4.1 | 0.001 |
| Movement * day | 6.6 | 2.2 | 3, 1058 | 5.4 | 0.001 |
| Mini-block * day | 3.9 | 0.3 | 15, 1058 | 0.6 | 0.84 |
| Movement * mini-block * day | 2.7 | 0.2 | 15, 1058 | 0.4 | 0.96 |

**TABLE 2.** STATISTICAL RESULTS FOR Δ_THRESHOLDS_.

|  | *SS* | *MS* | *dfs* | *F* | *p* |
| --- | --- | --- | --- | --- | --- |
| Mini-block | 16.2 | 3.2 | 5, 528 | 4.9 | < 0.001 |
| Day | 13.1 | 4.4 | 3, 528 | 6.6 | < 0.001 |
| Mini-block * day | 5.4 | 0.4 | 15, 528 | 0.6 | 0.92 |
| Movement speed | 0.1 | 0.1 | 15, 540 | 0.1 | 0.84 |

Our second aim was to quantify the reliability of the tactile perception and suppression estimates across different measurement points (mini-blocks and days). For this, we used permutation-based split-half correlations (e.g., [41, 42]), with custom-written programs in MATLAB 2024b. This analysis assesses the stability of the obtained perceptual measures across different measurement points, by comparing the consistency of the averaged values from two randomly selected halves. Specifically, for each variable (resting threshold, moving threshold, Δthreshold), we analyzed all 24 measurement values (6 mini-blocks x 4 days) for each of the 24 participants. The dataset of the complete data for each variable (6 mini-blocks * 4 sessions for all participants) was randomly split into two halves, with each half containing 12 randomly selected measurement values. For each half, we computed the average value for each participant based on the 12 observations and then calculated the Pearson correlation coefficient between these averaged values from the two halves. This procedure was bootstrapped 100.000 times, with new random splits generated in each iteration to minimize potential biases from specific divisions. The final reliability estimate for each of the three variables was determined by averaging the correlation coefficients.

Finally, our third aim was to relate the quantified temporal changes obtained from this experiment to previous findings of tactile suppression and assess whether tactile suppression may be driven by temporal changes in tactile sensitivity throughout an experiment. To address this, we reviewed the literature and obtained a weighted average of the tactile suppression effect size. We then compared the effect of any temporal changes observed in the present experiment to this tactile suppression effect. Details about this analysis and results are provided in the *Supplementary Material*.

## III. Results

We first tested if tactile detection is suppressed during movement both within and across days. As expected, detection thresholds were higher during movement than resting conditions, indicating tactile suppression (black lines and data points in Figure 2). Our main interest was in identifying changes in tactile detection thresholds across multiple measurement points over the span of a month. Main effects of mini-blocks and days were significant, but because these effects depended on the movement condition we will focus on the respective interactions (Table 1).

**Fig. 2.**
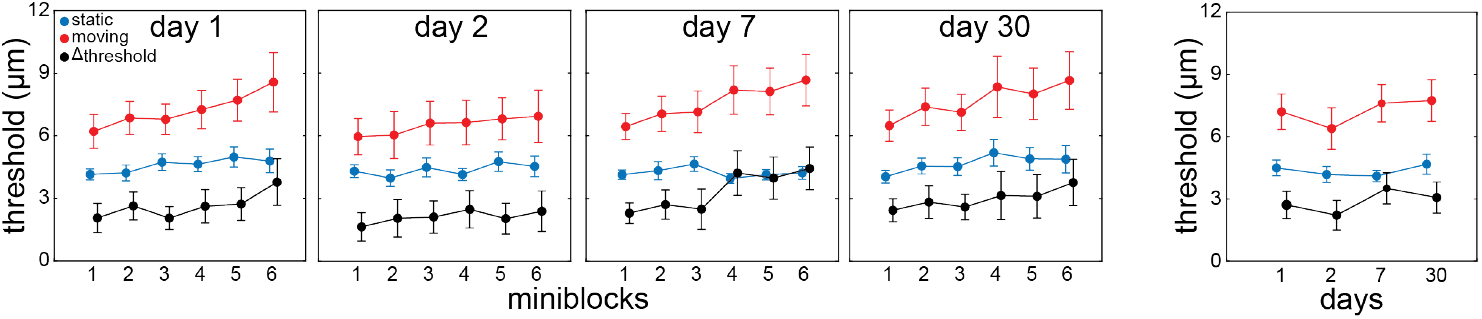
Tactile perception within and across days. Tactile detection thresholds during resting (blue) and moving (red) trials, as well as their relative difference (suppression; black), separately per mini-block and day (first four panels), and across days (last panel). Symbols represent averages across participants with error bars denoting the standard error of the mean.

Planned post-hoc comparisons exploring the movement by mini-block interaction revealed that detectionthresholds during the moving condition were lower in mini-block 1 relative to mini-blocks 4, 5, and 6 (all β > 0.44, all t > 4.84, all *d* > 0.30, all p < 0.001), as well as in mini-blocks 2 and 3 relative to mini-block 6 (both β > 0.43, both t > 4.68, both *d* > 0.29, both p < 0.001; red data in Figure 2). No differences between mini-blocks were found in the resting condition (blue data in Figure 2). These results suggest poorer perception with more exposure to the mini-blocks but only in trials involving movement. Planned post-hoc comparisons exploring the movement by day interaction revealed lower thresholds in the moving condition in day 2 relative to days 7 and 30 (both β > 0.37, both t > 4.92, both *d* > 0.30, both p < 0.001; Figure 2). Again, no differences across days were found in the resting condition. Together, the results demonstrate that detection thresholds can change both within and across days but only when the stimulated finger is moving. However, these effects are not systematic.

The second LMM focused on the tactile suppression effect (Δthresholds) and tested if tactile suppression changes within and across days while accounting for movement speed (e.g., 37, 43; but see also [44, 45]). There were main effects of mini-block and day on Δthresholds (see black lines and data points in Figure 2 and Table 2). Post-hoc pairwise comparisons on the effect of mini-block revealed significantly lower Δthresholds in mini-blocks 1 and 3 relative to mini-block 6 (both β > 0.43, both t > 3.63, both *d* > 0.32, both p <= 0.003). Likewise, post-hoc pairwise comparisons on the effect of day revealed significantly lower Δthresholds in day 2 relative to days 7 and 30 (both β > 0.28, both t > 2.95, both *d* > 0.26, both p <= 0.02; Figure 2). These results indicate that tactile suppression can change within short and longer time periods, but again, this is not systematic.

The second aim of our study was to examine whether tactile perception and suppression estimates were reliable across different measurement points. Split-half correlations showed strong consistency between the separate halves, with average Pearson correlation coefficients of 0.96 ± 0.01 for thresholds in static trials, 0.98 ± 0.01 for thresholds in movement trials, and 0.96 ± 0.01 for Δthresholds. The distributions of the bootstrapped correlation coefficients are shown in Figure 3.

**Fig. 3.**
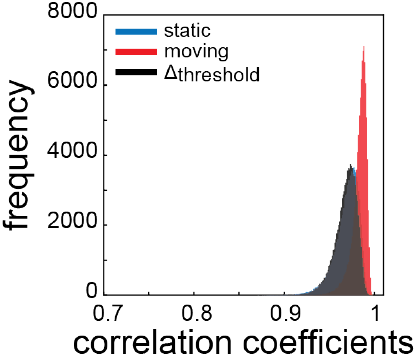
Split-half correlation coefficients. Distribution of Pearson correlation coefficients for split-half correlations, separately for thresholds estimated based on static trials (blue), movement trials (red), and Δthresholds (black).

## IV. Discussion

In this study, we examined whether tactile perception on a resting and a moving finger, as well as movement-related tactile suppression change with exposure to a tactile task. We addressed three research questions. First, do estimates of tactile perception and suppression change over time? Second, how reliable are the tactile perception and suppression estimates over time? Third, could any temporal changes in tactile perception over time explain the phenomenon of tactile suppression? To answer these questions, we asked participants to detect tactile stimuli on their right index finger, while this finger was resting or moving. These detection tasks were performed in ∼60-minute sessions that were repeated across four days within a period of one month. Our results demonstrate that tactile perception during rest is robust within the ∼60-minutes time window of each session. Likewise, we show generally robust tactile perception during rest also across days. In contrast, tactile perception during movement changed with more exposure to the task both within and across days. However, this effect was not always present, and thus not systematic. In addition, split-half correlations demonstrate robust estimates of tactile perception thresholds and associated tactile suppression over measurement periods.

Tactile detection thresholds on a resting finger were highly robust both within and across days. One might have expected improved detection with longer exposure to the experimental task and the tactile stimulation, for instance due to increased familiarity with the task and stimuli. Several studies have shown that repeated exposure to a task can cause a learning cascade of perceptual skills [26, 46, 47]. For instance, discrimination of auditory frequencies and visual motion directions can improve over seven consecutive days with repeated exposure to the task [27, 28], although such effects may require a minimum number of trials [27]. We have no evidence that tactile perception during rest improves with more exposure to the task and the chosen tactile stimuli. However, it is important to note that the tactile sensitivity of our participants during resting trials was very high, as they felt the weakest possible physical stimulus, making it impossible to observe any possible improvement over time. This is not surprising given the chosen frequency of our vibrotactile stimuli (250 Hz) which is known to be well detectable in humans (e.g., [48]). Such stimuli primarily activate the FAII rapidly adapting receptors that have large receptive fields [49, 50], which can, in turn, make the vibrotactile probes particularly salient. Although this might be considered a limitation when examining for possible improvements in resting tactile sensitivity over time, it is important to put this methodological decision in the context of the interests of this study, which sought to relate the current findings to existing evidence on tactile suppression that used probing stimuli of similar properties (i.e. frequency, duration, site of stimulation; [22, 30, 33]). One might have also expected that systematic stimulation of the tactile receptors on the index finger could lead to sensory adaptation and poorer sensitivity over time [21, 29]. Our data lend no evidence for this expectation. The mechanoreceptor properties that encode discriminative aspects of vibratory stimuli may make such adaptation unlikely, as the high frequency vibrations used in our experiment activate mechanoreceptors that adapt quickly in response to stimuli, primarily those moving across the skin [51, 52]. It can be argued that the fast-adapting properties of activated units contributed to the lack of adaptation in the rest condition. The robustness of resting tactile detection thresholds within and across days suggests that exposure to tactile stimuli does not have any worthwhile impact on tactile detection.

Tactile detection thresholds on a finger that was moving showed some change with more exposure to the task. However, these changes were not always observable and did not follow any consistent or systematic patterns. For instance, although detection thresholds during movement increased in some instances during the ∼60 minutes of the measurement sessions, the differences were not systematic across all mini-blocks. In addition, detection thresholds during movement were higher in the third (day 7) and fourth (day 30) relative to the second session (day 2), but there were no systematic differences with respect to the first day. We cannot exclude possible interactions between learning and adaptation effects. Specifically, it might be that a possible deterioration of perception evolved over time as a result of adaptation, but that participants also became better over time in detecting the stimuli in the moving trials, with these two effects eventually cancelling each other out. Together, these suggest that detection thresholds during movement can change with more exposure to the task, but the origin of these changes remains unclear. If the changes in thresholds were simply due to more exposure to the task, one might have expected the lowest detection thresholds to occur on the first measurement day. Yet, there was no difference between the first and any other day.

Tactile suppression was evident in all mini-blocks and sessions, in line with an abundance of studies showing poorer sensitivity to tactile stimuli presented around movement compared to rest (e.g., [7-12]). The strength of suppression was, to some extent, variable within days as it was larger in the last mini-block compared to the first and third. Although this reflects increased suppression with more exposure to the task within a single session, it should be interpreted with caution because we did not find any other systematic differences within single sessions. Likewise, although we found that tactile suppression changed across days, this change was neither linear nor systematic. Indeed, suppression was stronger in measurement days 7 and 30 relative to day 2, but not relative to day 1. Therefore, although exposure to the tactile task could lead to increased tactile suppression, this explanation is not enough to explain the observed pattern of results.

The reasons for the increased detection thresholds during the moving condition, as well as tactile suppression, on measurement days 7 and 30 relative to measurement day 2 remain unclear. Because the reported differences within and across days were found only when moving, kinematic factors might have influenced the related results. For instance, higher movement speeds can influence the magnitude of tactile suppression [37, 42]. This is the reason that we included movement speed as a covariate in our analyses. Another factor that could influence tactile perception is the time of stimulus presentation relative to movement onset [7, 53]. However, our stimuli were always presented between 10 and 80 ms (on average at 45 ms) after the onset of the movement, so this factor is unlikely to have driven temporal fluctuations. Therefore, the modulation on days 7 and 30 relative to day 2 may stem from unknown artifacts that could make the estimation of detection thresholds during movement less accurate. These findings may thus require further investigation, possibly by considering additional intervals when tactile perception is measured.

A second aim of our study was to examine the reliability of tactile perception estimates during resting and movement, and of the resulting tactile suppression. To assess this, we conducted a permutation-based split-half correlations analysis, which is used to evaluate the consistency in the measurements across different subsets of data [40, 41]. The results demonstrate that detection threshold and suppression estimates were remarkably robust across measurement points, with Pearson correlation coefficients consistently being above 0.9. This pronounced reliability highlights the stability of the calculated threshold estimates within and across sessions, providing confidence in the reproducibility of the observed effects.

Finally, the third aim of our study was to explore whether tactile suppression effects, as reported in the literature, may be caused by changes in tactile sensitivity over time (see *Supplementary Material*). Although it has been widely demonstrated that tactile suppression is modulated by various factors, such as movement speed [37], stimulation site [31], task demands [19], and the reliance on feedback and feedforward signals [23], one cannot exclude the possibility that tactile sensitivity changes throughout an experiment, such that thresholds in movement trials are systematically affected by temporal factors rather than by sensorimotor processes. For example, participants may experience fatigue, heightened anticipation, or adaptation to repetitive tactile stimuli over time, leading to altered thresholds that could confound suppression effects. Alternatively, differences in task engagement across rest and movement trials might also influence sensitivity, creating an apparent suppression effect that is not purely dependent on sensorimotor processes. To examine this, we evaluated whether the changes in tactile thresholds over time were smaller than the tactile suppression effects observed in the literature (see *Supplementary Material* for details on Analysis and Results). This is also the reason for using 250 Hz vibrotactile stimuli on the finger, which allowed us to match the methods of most previous studies on suppression. Thus, if suppression changes over time, the effect of exposure to the task would be equivalent to the suppression effect reported in the literature, independently of the performed movement. The related tests revealed that tactile suppression, as reported in the reviewed literature, cannot arise just from temporal changes in tactile sensitivity. Please note that the suppression effect size when considering only studies with detection of vibrotactile probes on the finger, similar to the experiment presented here, is even larger (please refer to the *Supplementary Material*). Still, we cannot exclude the possibility that temporal changes in perception may influence threshold estimates, but our results reflect that any temporal changes in tactile perception are not systematic. In short, the observed temporal changes in tactile perception cannot account for the tactile suppression described in previous studies.

We conclude that tactile perception during resting is robust both within and across days. Although tactile perception during movement and tactile suppression may change within and across measurement sessions, this change is not systematic. Measures of tactile perception and suppression are highly robust within and across measurement sessions, and any possible temporal variability cannot explain the movement-related tactile suppression effects reported in the literature.

**Dimitris Voudouris** received his PhD in human visuomotor control from the Vrije University Amsterdam in 2013. He continued at the same institute as a postdoctoral researcher, examining eye movements in grasping. Since 2014 he works at the Justus Liebig University Giessen, Germany. His research interests include tactile perception and gaze allocation during arm and whole-body movements.

**Petros Georgiadis** completed his master’s degree in Experimental Psychology at Justus Liebig University Giessen (2022–2024) and is currently pursuing a PhD in Biology at York University in Toronto, Canada. His research explores perception and action, focusing on the visual guidance of movement and brain-machine interface technologies.

**Katja Fiehler** performed her doctoral studies at the Max Planck Institute for Human Cognitive and Brain Sciences in Leipzig, Germany and received her doctoral degree from the University Leipzig, Germany in 2002. She continued her work on the interaction of somatosensory perception and action at the Philipps University in Marburg, Germany before she became full professor in Experimental Psychology at the Justus Liebig University Giessen, Germany. Her research focusses on human sensorimotor predictions and spatial coding for action in naturalistic environments.

**Belkis Ezgi Arikan** holds a PhD in Cognitive Neuroscience from Philipps University Marburg. From 2019 to 2024, she was a postdoctoral researcher at the Justus Liebig University Giessen. She is currently working at the Department of Neurology at University Hospital Essen. Her research interests involve action-perception coupling, prediction, time perception, multisensory processes and sense of agency. To investigate these, she uses behavioral (psychophysics) and neuroimaging (fMRI) methods.

