## Supplemental Material for "Reliability of tactile perception and suppression measurements"

for

One of our aims was to relate the findings of the present study to those from previous work and examine whether the previously reported suppression effects may arise as a byproduct of possible changes in tactile perception over time. In our study, this is reflected in the multiple measurement points spanning several days. To address this, we first quantified the effect of movement on tactile perception based on the literature (i.e., effect size of movement-related tactile suppression), and then tested whether the observed differences between the earliest and latest mini-blocks of our present study, as well as those between the first and the last days were within the margins of the effect size reported in the literature. To estimate the reported effect size, we reviewed the literature and found 41 studies on movement-related tactile suppression. Out of those, we considered 26 studies [8-10, 13, 14, 17, 18, 20, 22-25, 33, 42, 43, 54-63] that reported effect sizes in a direct (e.g., *Cohen's d*) or indirect manner (e.g., means, standard deviations and sample sizes). In case a study reported more than one related experiment, we used the smallest effect size across those experiments, which provides us with the most conservative effect. We calculated the Cohen's *d* from those studies, which we weighted based on the corresponding sample size (i.e., [64]), resulting in a true effect size of 1.28 (please see Table below for details). The studies included in this analysis involved mainly tactile detection tasks, various types of movements, and different tactile stimuli. When focusing only on studies conducted by our group, which included only tactile detection tasks with identical stimuli to those used in the present study (250 Hz vibrations of 50 ms duration and similar intensities), we observed that the weighted effect size was slightly higher than that observed in other studies (1.32 vs 1.23), but also larger than the grand weighted average effect size (1.29), which highlights that we use the most conservative effect size. We then used two independent one-sided t-tests to examine whether the observed differences in tactile perception and suppression between the first and last mini-blocks as well as between the first and last measurement days were within the margins of the true effect size (i.e. [-1.29 to 1.29]; equivalence testing; [65]). These equivalence tests, if statistically significant, will inform about the temporal effect observed in the current experiment being smaller than the suppression effect reported in the literature, and, thus, tactile suppression being unlikely to be driven by temporal changes in tactile sensitivity. To this end, we first averaged the detection thresholds in each of the static and moving conditions, as well as the tactile suppression indices, across the earliest and latest mini-blocks of each measurement day, obtaining an *early* and a *late* detection threshold for each of the resting and moving conditions, as well as an *early* and a *late* tactile suppression index. We also examined whether the observed differences in tactile perception and suppression between the first and last measurement days were within the margins of the true effect size. To this end, we fitted new psychometric functions for each measurement day, separately for each (resting and moving) condition and per participant, by merging the responses across all mini-blocks of the respective condition and the respective day. This can result in more reliable fits while still allowing us to obtain detection thresholds for the resting and moving conditions, as well as the resulting tactile suppression indices. The equivalence tests were conducted in Jamovi 2.5.4 [66] using the TOSTER package.

We examined whether the observed effect between the first and the last mini-blocks as well as between the first and last measurement day was smaller than the suppression effect size as we calculated it based on the literature (see above). The two one-sided t-tests of the equivalence testing procedure applied separately to the resting and moving thresholds, as well as to the *Δthresholds*, reveal that the observed differences between the first and last mini- blocks within measurement days were negligible to explain the phenomenon of tactile suppression. Indeed, the observed effect sizes were within the margins of the true effect size for the resting detection thresholds (both $|t| > 4.85$, both $p < 0.001$), the moving detection thresholds (both $|t| > 4.15$, both $p < 0.001$), and the *Δthresholds* (both $|t| > 4.37$, both $p < 0.001$). Likewise, the differences between the first and the last measurement days were smaller than the suppression effect size as reported in the reviewed literature. Specifically, the observed effect sizes of the differences between the first and last measurement day were within the margins of the suppression effect size for the resting detection thresholds (both $|t| > 5.54$, both $p < 0.001$), the moving detection thresholds (both $|t| > 4.99$, both $p < 0.001$), and the *Δthresholds* (both $|t| > 5.41$, both $p < 0.001$). We can thus argue that possible differences in tactile perception and suppression within and across measurement days cannot be the main reason for the tactile suppression effect as these are described in the reviewed literature (see table below).

| no. | entry in reference list | paper | movement | tactile task | stimulus at | type of stimulus | Cohen's d | N | weighted value |
|---|---|---|---|---|---|---|---|---|---|
| 1 | 8 | Juravle et al., 2018, Sci Rep | grasping | detection | finger | electrical | 2.50 | 20 | 50 |
| 2 | 9 | Manzone et al., 2018, BBR | grasping | detection | finger | electrical | 0.75 | 11 | 8.25 |
| 3 | 10 | Voudouris & Fiehler, 2017, JEP: HPP | reaching | detection | finger | vibration | 1.68 | 18 | 30.24 |
| 4 | 13 | Fuehrer et al., 2022, PNAS | stroking | detection | finger | vibration | 1.11 | 32 | 35.52 |
| 5 | 14 | Wachsmann et al., 2023, biorxiv | standing | detection | lower leg | vibration | 0.72 | 10 | 7.2 |
| 6 | 17 | Arikan et al., 2024, iScience | wrist rotation | detection | finger | vibration | 1.90 | 37 | 70.3 |
| 7 | 18 | Chapman & Beauchamp, 2006, JNP | finger abduction | detection | finger | electrical | 1.30 | 8 | 10.4 |
| 8 | 20 | Gertz et al., 2018, PLoS ONE | reaching | detection | finger | vibration | 1.85 | 13 | 24.05 |
| 9 | 22 | Broda et al., 2020, Sci Rep | grasping | detection | finger | vibration | 1.63 | 12 | 19.56 |
| 10 | 23 | Voudouris et al., 2019 Sci Rep | grasping | detection | finger | vibration | 1.04 | 20 | 20.8 |
| 11 | 24 | Kilteni & Ehrsson, 2022, iScience | force application | discrimination | finger | force | 0.50 | 24 | 12 |
| 12 | 25 | Arikan et al., 2021, Neuroimage | reaching | detection | finger | vibration | 3.70 | 12 | 44.4 |
| 13 | 30 | Voudouris & Fiehler, 2022, HMS | grasping | detection | finger | vibration | 0.90 | 20 | 18 |
| 14 | 33 | Gertz et al., 2017, JNP | reaching | detection | finger | vibration | 2.32 | 12 | 27.84 |
| 15 | 43 | Angel & Malenka, 1982, Exp Neurol | tracking | detection | finger | electrical | 1.05 | 8 | 8.4 |
| 16 | 44 | Voudouris & Fiehler, 2021, Sci Rep | reaching | detection | finger | vibration | 1.09 | 16 | 17.44 |
| 17 | 54 | Beyvers et al., 2022, Front Neurosci | reaching | detection | finger | vibration | 0.42 | 26 | 10.92 |
| 18 | 55 | Beyvers et al., 2023, Sci Rep | grasping | detection | finger | vibration | 1.44 | 24 | 34.56 |
| 19 | 56 | Juravle & Spence, 2011, EBR | juggling | detection | wrist | vibration | 1.58 | 10 | 15.8 |
| 20 | 57 | Juravle & Spence, 2012, EBR | catching | detection | wrist | vibration | 2.70 | 14 | 37.8 |
| 21 | 58 | Juravle et al., 2011, Acta Psychol | grasping | detection | finger | vibration | 1.32 | 12 | 15.84 |
| 22 | 59 | Juravle et al., 2010, BBR | grasping | discrimination | finger | vibration | 0.98 | 19 | 18.62 |
| 23 | 60 | Klever et al., 2019, JoV | reaching | detection | finger | vibration | 0.65 | 23 | 14.95 |
| 24 | 61 | Schuetz et al., 2022, Haptics | reaching | detection | palm | vibration | 0.77 | 18 | 13.86 |
| 25 | 62 | Thomas et al., 2022, Psych Sci | force application | discrimination | force | finger | 0.50 | 24 | 12 |
| 26 | 63 | van Hulle et al., 2013, Consci Cogn | back bending | detection | lower back | vibration | 0.98 | 12 | 11.76 |

| | | | |
|---|---|---|---|
| **weighted effect size** | | | **1.29** |
| weighted effect size (own group studies) | | | 1.32 |
| weighted effect size (studies from other groups) | | | 1.23 |
| weighted effect size (only studies with vibrotactile detection on the hand) | | | 1.38 |
